# Snake Venom Fluidic Properties and Design of Venom Mimics as Rheological Surrogates

**DOI:** 10.64898/2026.06.19.733472

**Authors:** Madison Forstner, Matthew L. Holding, Yiqun Li, Talia Y. Moore, Abdon Pena-Francesch

## Abstract

Snake venom composition and its contribution to toxic effects has been heavily researched, but there is a comparative lack of information on venom’s fluidic properties and their relationship with fang morphology during the envenomation process. Understanding how venom flows through a fang can shed light on bite site dynamics and potentially explain bite symptoms. In this article we first conduct a broad comparative test of the rheological properties of venom from thirteen snake species, including multiple viperid and elapid snake species, revealing a shear-thinning non-Newtonian flow behavior in all studied species. However, we have not observed strong phylogenetic signal in venom fluidic properties, suggesting that flow properties may vary independently of evolutionary relationships between snake species. Second, we demonstrate that snake venom’s fluidic properties can be modeled by other inexpensive, safe, and abundant shear-thinning surrogate fluids. We found that aqueous solutions of bovine serum albumin protein and xanthan gum are useful venom mimics, matching the rheological behavior of venoms from the studied snake species across a range of relevant shear rates. We further evaluated the performance of these snake venom mimics in a simulated venom delivery system, showing good and robust mimetic control of the flow properties as a function of applied pressure. By elucidating the fluidic properties of snake venom and providing a non-toxic, scalable surrogate fluid model to be used in further studies, we provide the biomedical, toxicology, evolutionary biology communities with a tool to study envenomation physics in an inexpensive and safe fashion. We suggest it is possible to design species-specific venom mimics that facilitate research on the biomechanics and fluid dynamics of venom delivery *via* snake bites, and inform the design of bioinspired puncture and injection devices.

## Introduction

In 2017, snakebite envenoming was recognized by the World Health Organization (WHO) as a Category A Neglected Tropical Disease. Snakebites are far-reaching with 5.4 million bites per year worldwide resulting in 1.8 to 2.7 million cases of envenoming, 137,000 deaths, and three times as many amputations and other permanent disabilities annually^1,2^. Snakebite nonetheless remains a neglected public health issue due to the low production of antivenoms, lack of health and distribution infrastructure, and poor data on the number and type of snake bites to provide fast, appropriate treatment. Snake venom composition (comprising proteins, enzymes, amines, lipids, carbohydrates, nucleosides, and metal ions) and the resulting toxic effects of venom have been the target of most contemporary research on snake envenomation^3^. Protein concentration in venom can be quite high (100-300 mg/mL)^4,5^, which is expected to affect rheological properties relevant to venom delivery. This suggests that snakes may be navigating an evolutionary tradeoff as higher protein concentrations enhance venom toxicity, but viscosity in ways that may hinder delivery.

However, there is a comparative lack of information on venom fluidic properties and their relationship with fang morphology during the envenomation process. For example, in front-fanged venomous snakes, venom is contained in a gland, which must be compressed to force the venom through a duct and into and through the hollow fang^6,7^. The functional morphology of venom injection (**Figure 1a**) dictates that venom must: (i) not flow out while stored in the venom gland, (ii) start flowing from the gland at a specific pressure threshold, (iii) flow through the fang microchannel or side-grooves rapidly, and (iv) flow and diffuse into the prey tissue. These requirements suggest adaptive viscosity behavior typically observed in shear-thinning non-Newtonian fluids^8^. As such, venom would remain highly viscous in the gland until the snake’s compressor muscle forces the venom out of the venom gland and into the primary venom duct, placing shear forces onto the venom inside and reducing its viscosity to enable its flow^9–11^.

**Figure 1.**
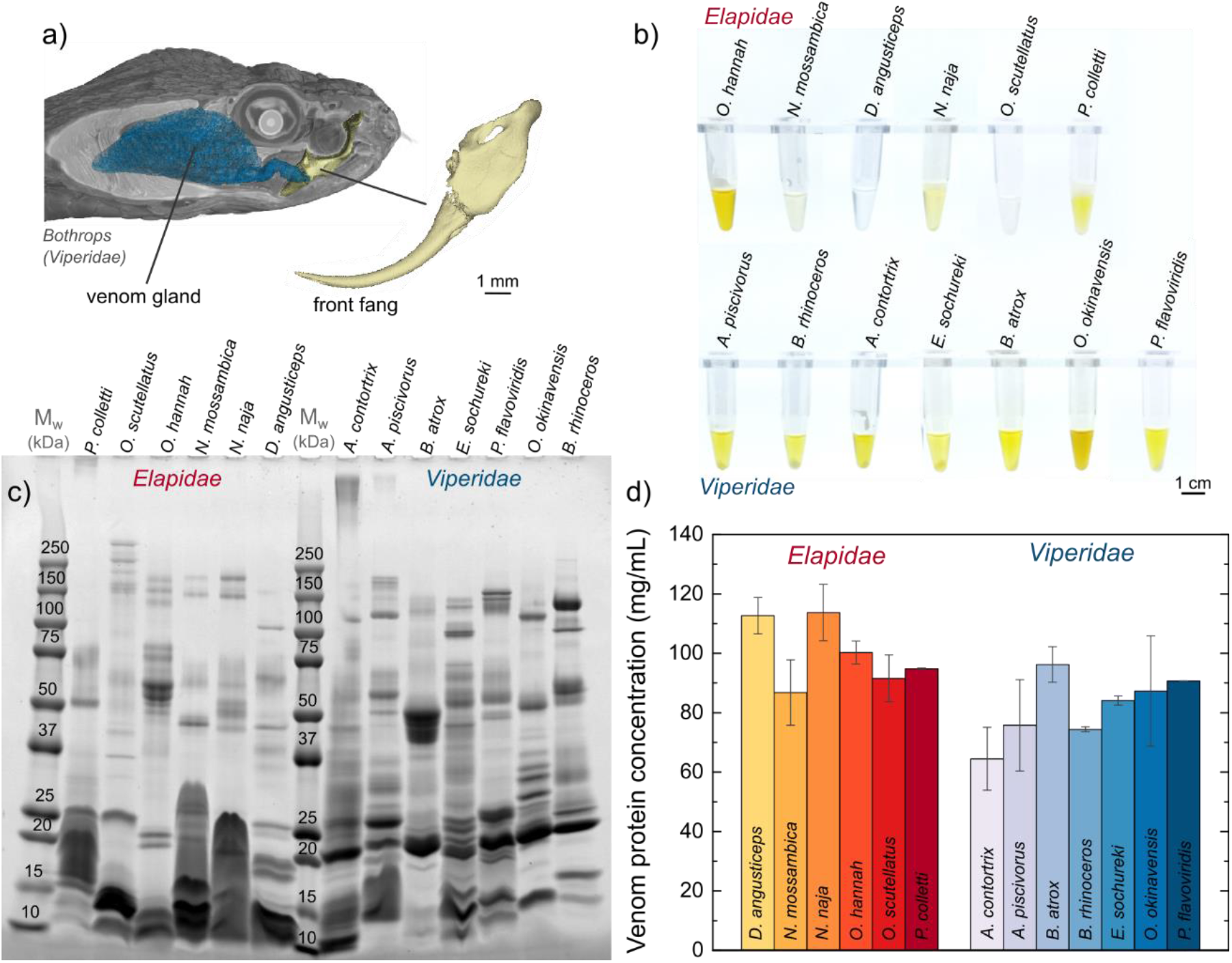
Snake venom proteins. **a)** Venom delivery system of a *Bothrops* (*Viperidae*) snake (CT scan), with the venom gland highlighted in blue and the front fang magnified. Adapted with permission from ref^24^. **b)** Venom samples from elapid and viper snakes after centrifugation, ranging from colorless to intense yellow. **c)** Molecular weight distribution of venom proteins of elapid and viper snakes, measured by gel electrophoresis, in the 10 - 250 kDa range. **d)** Protein concentration in venom of elapid (red) and viper (blue) snakes, measured by BCA.

The shear-thinning properties of venom were first shown in the rattlesnakes (Viperidae)^12^ and then the cobra family (Elapidae).^9^ The extensive natural variation in venom composition, venom concentration,^13^ snake size,^14^ and fang morphology^15^ raise the question that fluidic properties might be an important feature in venom evolution, and that fluidic properties of venom themselves may show adaptive variation. Moreover, variable fluidic properties among venoms could interact with venom composition to dictate the movement of venom into a wound and subsequent bite symptomology. To our knowledge, a single prior study explored this possibility, with a focus on spitting and non-spitting cobras^16^. Despite the special role of long-distance venom spitting in spitting cobra biology, the viscosity of venom at high shear rates (~10,000 s^−1^) did not differ between spitting and non-spitting cobra species, calling into question the nature of snake venom as a non-Newtonian liquid. Clearly, additional comparative research on multiple snake species over a broad range of shear rate conditions and with different venom compositions is needed to elucidate composition-property-function physical relationships that describe the fluidic behavior of venom.

Furthermore, studying the relationships between venom fluidic properties, the morphology of the venom delivery system, and the kinematics of snakebite would be facilitated by the identification of appropriate venom mimic fluids. Current venom extraction processes depend on live snakes, and thus venom can be costly to acquire, limited in availability, and of course dangerous. A mimic fluid model that can replicate the venom’s fluidic properties with inexpensive, safe-to-handle, non-toxic formulations would provide a robust tool for the research community to study envenomation physics at larger volumes and length scales.

Here, we describe a quantitative study of the rheological properties of venom from thirteen *Viperid* and *Elapid* snake species, using steady shear measurements over a wide range of conditions and systematically analyzing their shear-thinning behavior. We conducted a comparative analysis of venom fluidic key rheological parameters to test for differences between venom composition and phylogenetic signal. By mapping the variable viscosity as a function of shear rate for all the studies samples, we formulated a series of venom mimic fluids based on protein (bovine serum albumin, BSA) and polysaccharide (xanthan gum) aqueous solutions. With these mimic fluid models, we observed similar rheological behavior to snake venom from vipers and elapids, and we demonstrated switchable flow behavior in a pressure-driven direct ink write (DIW) nozzle 3D printer system as a simulated fang envenomation system.

## 2. Methodology

### Materials

Snake venom samples were purchased as lyophilized powder from the Kentucky Reptile Zoo (Slade, Kentucky, United States). Eight samples were selected from the *Viperidae* family and six from the *Elapidae* family. The snake species and abbreviated identifier are listed in **Table 1**. Bovin serum albumin (BSA, CAS 9048-46-8) protein was purchased from MilliporeSigma, and food-grade xanthan gum (UPC 850005727064) was purchased from Amazon.

**Table 1:** Snake species and identifiers.

| Common name | Latin name | ID |
| --- | --- | --- |
| Vipers |  |  |
| Copperhead | *Agkistrodon contortrix contortrix* | ACC |
| Cottonmouth | *Agkistrodon piscivorus* | AP |
| Brazilian Lancehead | *Bothrops atrox - Suriname* | BAT-S |
| Saw-scaled viper | *Echis sochureki* | ES |
| Okinawan pitviper | *Ovophis okinavensis* | OvO |
| Habu | *Protobothrops flavoviridis* | PF |
| Gaboon viper | *Bitis rhinoceros* | BR |
| Elapids |  |  |
| Green mamba | *Dendroaspis angusticeps* | DA |
| Mozambique spitting-cobra | *Naja mossambica* | NMo |
| Indian cobra | *Naja naja - Indian origin* | NNA-I |
| King cobra | *Ophiophagus hannah* | OH |
| Coastal taipan | *Oxyuranus scutellatus scutellatus* | OSS |
| Collett's black snake | *Pseudechis colletti* | PC |

### Preparation of snake venom solutions

We studied venom samples from eight snakes in the *Viperidae* family and six in the *Elapidae* family (**Table 1)**. Venom samples were milked from adult snakes and lyophilized at the reptile zoo facility, and were received as a dry white powder. Unfortunately, the initial venom volume that was milked was unknown, and therefore the initial concentration of the samples before lyophilization was unknown. Venom concentration can show high variability between specimens, and can be dependent on diet, specimen hydration, and enclosure conditions, and it can contain other solids besides venom proteins (cell debris, aggregates, etc.). To reliably compare species under relevant conditions, we reconstituted the venom samples to a solid lyophilized powder concentration of 150 mg/mL in deionized water. This falls within a venom concentration range observed in many species^16^. The venom samples were thoroughly mixed overnight to ensure that all components were dissolved, and then were centrifuged for 10 minutes at 10,000xG to separate any remaining solid fraction. The liquid fraction of the venom solution was isolated and analyzed for protein content using the Pierce BCA Protein Assay^17^. The purified venom samples were at −18 ℃ for later use.

### Venom protein content analysis

We further adjusted concentration of the protein supernatant based on the results of a Pierce BCA Protein Assay following manufacturer protocols. We loaded 20 µg of each sample into 15-well NuPAGE 4-20% Tris Glycine gels (Invitrogen). The gels were electrophoresed for 1 hour at 200V and the proteins were stained using Simply Blue Safe Stain (Invitrogen). The gel was imaged on a ChemiDoc MP Imaging System (BioRad) and relative abundance and molecular weight of the visible bands was quantitated using BioRad ImageLab v. 6.1.

### Rheological characterization

Venom solutions from all snake species were prepared at a concentration of 150 mg/mL. BSA solutions were prepared at concentrations of 5, 10, 20, 50, 100, 150, and 200 mg/mL. Xanthan gum solutions were prepared at concentrations of 0.2, 0.3, 0.4, 0.5, 0.6, 0.7, 0.8, 0.9, 1, 1.5, 2, 2.5, and 3 mg/mL. Each sample was mixed within a vibrational mixer and centrifuged for 5 minutes to remove impurities. A TA Instruments DHR30 rheometer was used to perform all viscosity characterization measurements. A stainless steel 2 deg, 20 mm cone and plate were utilized in steady-shear experiments (75 µL of sample volume) at a controlled temperature (20 ℃) under shear rate sweeps from 1000 to 0.1 s^−1^. Between each sample the top and bottom geometries were cleaned with isopropyl alcohol and water. When the experiment concluded, any tools in contact with venom samples were further sanitized with a 10% bleach solution to neutralize any lingering venom. BSA and xanthan gum solutions at low concentrations exhibited low viscosity, and were additionally characterized using a double gap concentric cylinder geometry (13 mL of sample volume), which is more sensitive to low viscosity solutions. The results from both cone and plate and double gap trials were combined.

### Comparative analyses

We tested for phylogenetic signal in high shear rate viscosity, crossover shear rate, and flow behavior index using the “phylosig” function in the phytools R package^18^. We obtained a time-calibrated phylogenetic tree of focal snake species using TimeTree 5^19^. We specified Blomberg’s K as the statistic of interest, and used 1000 randomizations to determine the statistical significance of the resulting K value.

### Simulated delivery of venom tests

An INKREDIBLE+ (Cellink, Sweden) 3D printer (connected to an external air compressor) was used to simulate venom delivery and dripping experiments perform the dripping test. 40% glycerol, 1 mg/mL xanthan gum, and 200 mg/mL BSA solutions were selected as reference fluids and venom mimics for validation. Venoms labeled NNA-I and BAT-S were selected and prepared as previously described for rheological tests. The solutions were extracted with a 3 mm luer-lock syringe and transferred to the printing cartridge. A 27G (200 μm) nozzle was used during the dripping test. For each solution, pressure was steadily increased from 0 kPa to 15 kPa with 1 kPa increments. The solution was allowed to flow for 10 seconds at each pressure step and the collected delivered solution was weighed.

## 3. Results and Discussion

### Characterization of snake venom

The venom samples studied all have a distinct visual appearance, with varying opacity and shade of yellow color (**Figure 1b**). Elapid samples showed a clear light shade of yellow, with the exception of *O. hannah* sample. Viperid samples shower a more intense, opaque yellow color, with a larger visible solid pellet after centrifugation. Venom composition in *Elapidae* and *Viperidae* snakes have been reported in extensive databases.^3^ While there is substantial diversity within species and subspecies, elapids have three-finger toxins (3FTxs, average 51%) and secreted phospholipases A2 (PLA2s, 27%) as major dominant constituents of the venom proteome, and viperids have snake venom metalloproteinases (SVMPs, 33%), PLA2 (21%), and snake venom serine proteases (SVSPs, 15%).^3^ 3FTxs (major constituent of elapid venom) are small proteins with a molecular weight <10 kDa (**Figure 1c**, likely comprising the observed dark, smeared bands on the bottom left of the gel image).^20,21^ Metalloproteinases (major component of viperid venom) are larger in size, and are typically found in viperids in three groups: P-I (20-30 kDa, single metalloprotease), P-II (30-60 kDa, which contain a metalloproteinase domain, a prodomain, and a disintegrin domain), and P-III (60-110 kDa, which contain a metalloproteinase domain, a prodomain, a disintegrin-like domain, and a cysteine-rich domain).^22,23^ Metalloproteinases and serine proteases are observed in viperid gel lanes densely localized in 20-25 kDa bands, 50 kDa bands, and in some samples in >75kDa bands. Despite these general trends, there is substantial diversity across species. For example, some venom samples, such as *P. colletti, A. contortrix*, and *A. piscivorus* (and other samples at a lower degree), exhibited large molecular weight (>250 kDa) stained regions at the top that did not go through the gel, suggesting the formation of large molecular weight protein aggregates. *O. hannah* and *B. atrox* are mainly composed of medium-sized proteins, while *P. colletti, N. mossambica, N. naja, D. angusticeps*, and *A. piscivorus* seem to be mainly composed of small proteins. *Elapidae* snakes exhibited concentrations in the 85-115 mg/mL range and *Viperidae* snakes in the 60-90 mg/mL range (**Figure 1d**).

### Venom rheological behavior

The snake venom samples exhibited an apparent non-Newtonian, shear-thinning behavior, with viscosity decreasing steadily with shear rate (**Figure 2**)^8,25^. This behavior is common in many biofluids^26^; at rest, proteins, polysaccharide, and other biomolecules form a network of intermolecular interactions that result in high viscosity. As the fluid is sheared, these interactions are disrupted and facilitate flow, thus reducing the viscosity. Over the range of shear rates studied here, we confirm the shear-thinning behavior previously observed in rattlesnake and cobra venom,^16,27,28^ showing lower viscosity at high shear rates. These non-Newtonian flow properties would benefit the mechanisms of envenomation of the studied species, where high viscosity at low shear rates would help to store the venom in the glands and low viscosity at high shear rates would help flow during delivery.

**Figure 2.**
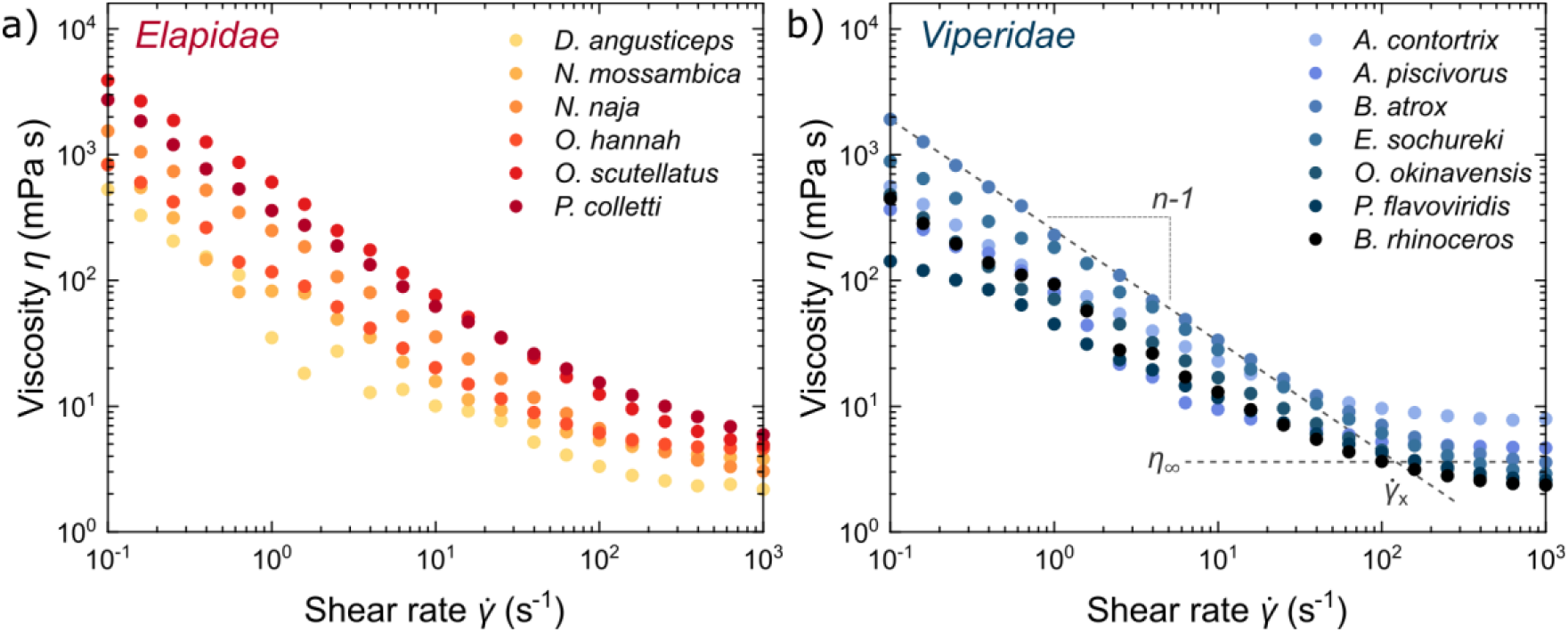
Rheological characterization of snake venom. **a)** Steady shear flow measurements of elapid snake venom. **b)** Steady shear flow measurements of viper snake venom. Viscosity as a function of shear rate shows a shear-thinning behavior, characterized by *n* (flow behavior index), *η*_∞_ (high-shear viscosity), 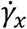 rate), n-1 represents the slope of the shear-thinning region.

Despite their general shear-thinning behavior, there are some apparent differences between elapids (**Figure 2a**) and viperids (**Figure 2b**), and between species within the same family. We used three characteristic parameters to compare the flow behavior of each venom sample and later measured the phylogenetic signal of each parameter across a phylogenetic tree (**Figure 3**).

**Figure 3.**
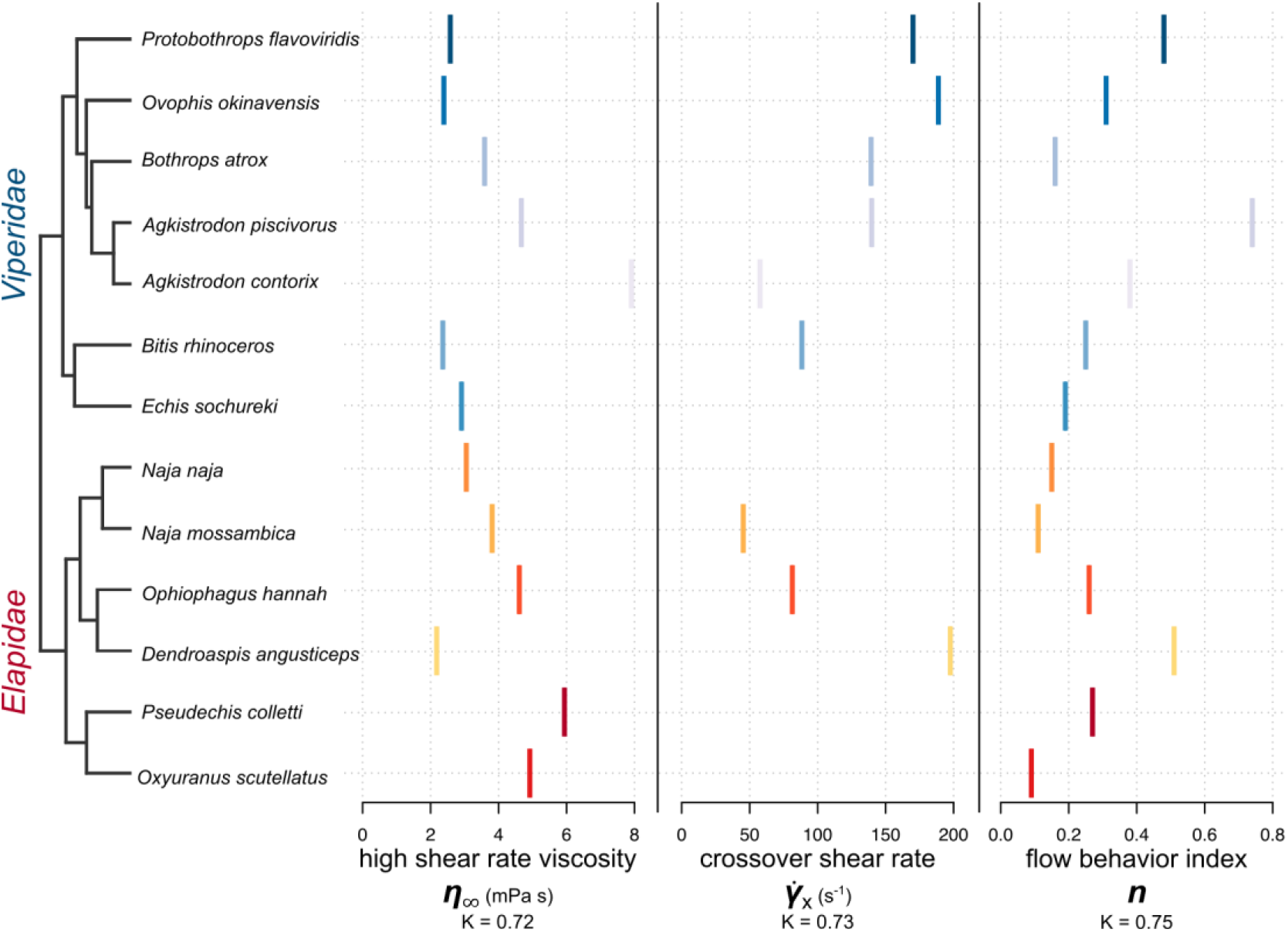
Analysis of the flow behavior of snake venom. Viper (blue) and elapid (red) snake venom characterized by *η*_∞_ (high-shear viscosity), 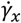 (crossover shear rate), *n* (flow behavior index), arranged by closest evolutionary neighbor in a phylogenetic tree. Bloomberg’s K did not show statistical significance among the studied species and data.

First, we modeled the viscosity using the power-law Ostwald–de Waele relationship to quantify the non-Newtonian behavior.^25^ This power-law relationship describes the viscosity as 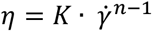, where *K* is a constant based on the fluid’s intrinsic viscosity at a shear rate of 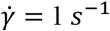 and *n* is the flow behavior index characteristic of the fluid. Newtonian fluids have a flow behavior index of *n* = 1 (shear rate-independent viscosity), while shear-thinning non-Newtonian fluids will present a flow behavior index of *n* < 1. In all samples, we observed the shear-thinning region in the 0.1 - 100 s^−1^ range. Some species had unusually high viscosities. For example, *A. contrix* and *A. piscivorous* were the two viperids with highest *η*_∞_, while *P. colleti* and *O. scutellatus* were the most viscous elapid venoms. These species also exhibited high molecular weight protein content or protein aggregates in the electrophoresis gel, which may be responsible for their higher viscosities.

Second, at high shear rates, the viscosity reached a rate-independent plateau at *η*_∞_, which represents the Newtonian flow behavior at “infinite” shear rate. Venom samples from both viperid and elapid families exhibited high shear rate viscosities *η*_∞_ between 2 and 6 mPa·s. These viscosities are lower than those reported for other families like spitting cobras^16^, possibly due to differences in how the venom samples were prepared. For example, while other reports have used freshly-extracted venom, we used venom that had been previously lyophilized and then clarified by centrifugation to obtain precipitate-free venom.

And last, we measured the crossover point between the shear-thinning and Newtonian regimes. The crossover point 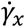 is the shear rate at which the fit line from the power law model intercepts the high shear viscosity plateau. While 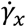 was easier to observe in viperids, it was not observed in a few elapids, as the Newtonian plateau was not reached in the studied shear rate range, indicating a stronger shear-thinning behavior over a wider range.

Overall, we did not find evidence for phylogenetic signal in the measured venom viscosities, suggesting that rheological parameters of venom vary independently of relationships between snake species. A narrow range of parameter values would indicate strong stabilizing selection on venom viscosity, potentially due to a biomechanical constraint on delivery, but we observed the opposite. The relatively large range and low phylogenetic signal in parameter values suggests that variation in venom viscosity may be the neutral consequence of selection on specific toxins within the venoms.

### Design and characterization of venom mimics

Having characterized all the venom samples and mapped out their characteristic flow parameters, we set out to design a mimetic venom surrogate. First, we explored bovine serum albumin (BSA) protein solutions as a candidate to formulate the venom mimics. BSA is a monomeric carrier protein (66.5 kDa) found in the blood serum of cows that is commonly used as model protein in many branches of biochemistry. Due to the ubiquity of BSA, there is extensive documentation on its rheological properties and as a shear-thinning biofluid^29–31^. We remeasured those here to standardize the comparison based on our own experimental approach, spanning concentrations up to 200 mg/mL (**Figure 4a**). Overall, BSA showed shear-thinning behavior with viscosity decreasing steadily with shear rate in the 0.1 to 10 s^−1^, following the same trend in all protein concentrations. However, at higher shear-rates (~100 s^−1^), the viscosity transitioned into a Newtonian regime with a plateau at *η*_∞_ (ranging from 1 to 4 mPa·s). The high shear rate viscosity *η*_∞_ increased linearly with protein concentration, similar to previous reports (**Figure 4a inset**).^31^ However, the viscosity range accessible through the *venom mimic I* is more limited than the viscosity range observed in snake venom. Even at higher protein concentrations, the *mimic I* viscosities are lower across the studied shear rates with only partial overlap in the high shear rate Newtonian region, and is only able to meet the high shear viscosity for eight species of snake venom out of thirteen (**Figure 4b** and **4c**). Therefore, due to the mismatch in rheological properties, we moved on to other mimic formulation candidates.

**Figure 4.**
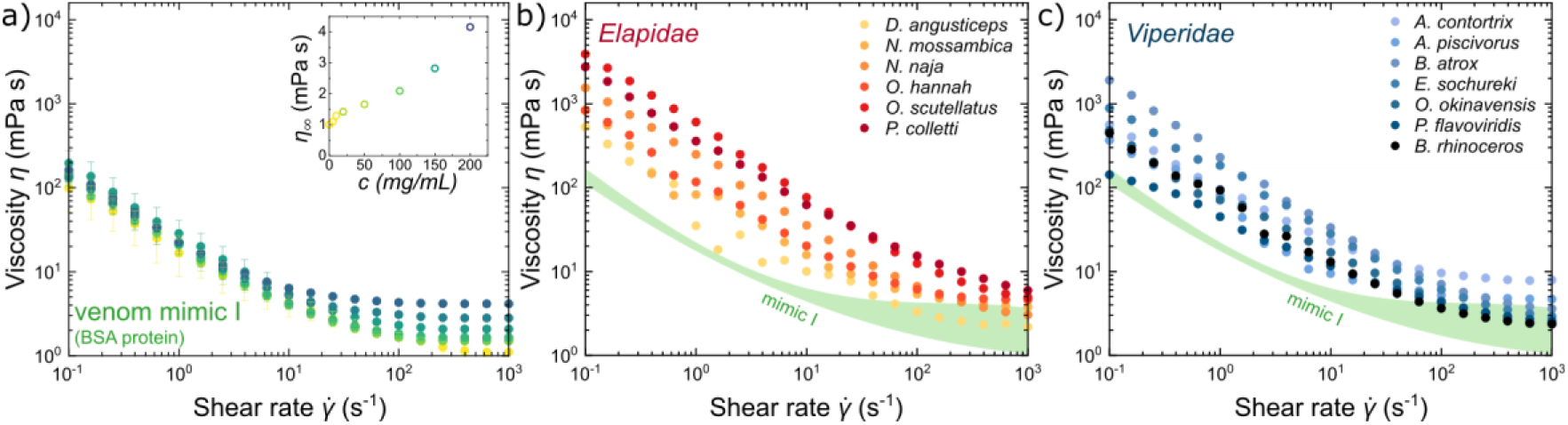
Rheological behavior of protein-based venom mimic. **a)** Steady shear flow measurements of BSA protein solutions (venom mimic I). Inset shows high-shear viscosity *η*_∞_ as a function of concentration. **b)** Elapid snake venom and **c)** viperid snake venom flow behavior overlaid on the accessible flow range of venom mimic I (in green).

As an alternative formulation, the *venom mimic II* was composed of xanthan gum aqueous solutions (**Figure 5a**). Xanthan gum is a pentasaccharide commonly used as a food additive and thickening agent, and its non-Newtonian shear-thinning properties have been extensively studied.^32–35^ *Venom mimic II* exhibits a shear-thinning behavior across all the studied shear rates, and is able to achieve a large range of viscosities at low concentrations. *Venom mimmic II* exhibits high shear rate viscosities *η*_∞_ from 2 to 12 mPa·s at concentrations from 0.1 to 3 mg/mL (**Figure 5a inset**), which encompass the full range of viscosities observed in real snake venom. Compared to the protein-based *mimic I*, the *venom mimic II* system requires much smaller concentrations to achieve the desired non-Newtonian behavior in xanthan gum formulations, and is able to match the rheological profiles of all thirteen venom samples (including both viperids and elapids) (**Figure 5b** and **5c**). Furthermore, xanthan gum is commercially available and has a relatively low cost due to its extensive use as food additive, making *venom mimic II* formulations easy to scale-up for large volume tests and further experimentation.

**Figure 5.**
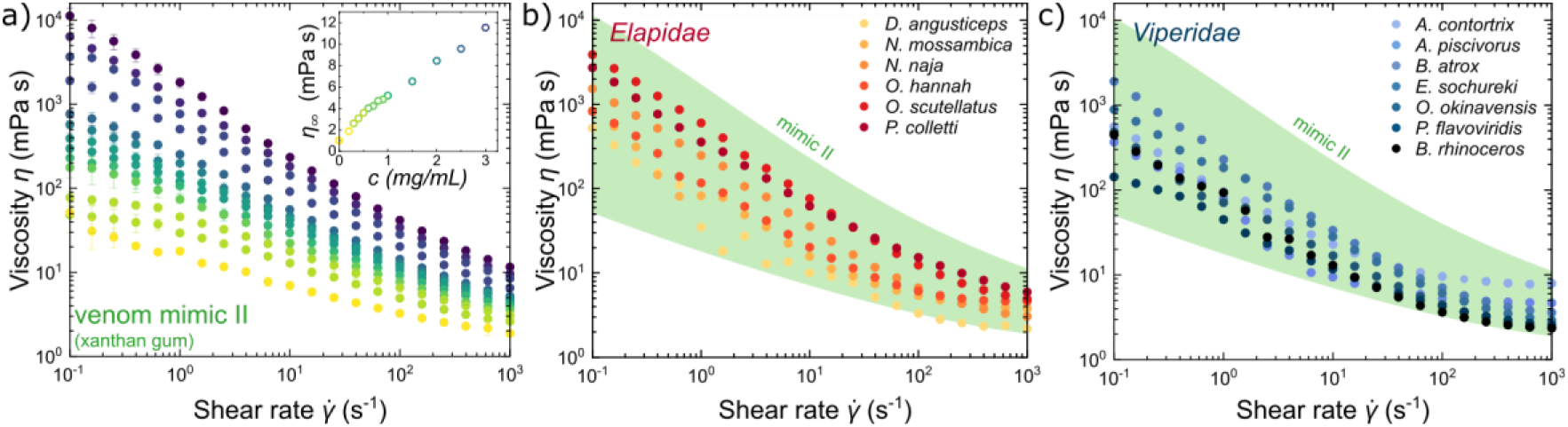
Rheological behavior of sugar-based venom mimic. **a)** Steady shear flow measurements of xanthan gum solutions (venom mimic II). Inset shows high-shear viscosity *η*_∞_ as a function of concentration. **b)** Elapid snake venom and **c)** viperid snake venom flow behavior overlaid on the accessible flow range of venom mimic II (in green).

### Simulated delivery of venom and mimics

After characterizing the rheological properties of venom from 13 snake species and developing two venom mimic models, we simulated the delivery of venom with real venom and mimic samples to evaluate their flow behavior under relevant flow conditions. We adapted a commercial direct ink writing 3D printer using syringe cartridges, extrusion nozzles 200 μm (G27) in diameter, and a satellite pump and pneumatic circuit to provide a versatile benchmarking platform and simulate venom delivery with controllable pressure. Venom samples were loaded in the cartridges and pressure was applied from 0 to 15 kPa in step functions of 1 kPa increment (**Figure 6a**). The amount of extruded fluid under each pressure condition was measured and normalized by delivery time to provide a flow estimate and compare venom formulations. Due to the low availability of real venom samples and the volumes required for testing, we were able to measure only two venom samples (one elapid *N. naja* and one viperid *B. atrox*) at limited pressure and flow conditions. However, it was enough to observe a representative flow trend. At low pressures (3 kPa) the real venom samples did not flow at all, and only at 7 kPa pressures we observed flow out of the nozzle. After the initial no-flow regime at low pressures, both venom samples showed a linear flow dependence with increasing pressure, which was expected in a shear-thinning fluid. These measured pressures to achieve flow in our simulated 3D-printer-based system are lower than others reported for rattlesnakes and cobras (within the 50 kPa range)^6,16^, probably due to the smaller size of snake fang venom channels (few micrometers in diameter). For reference, a Newtonian fluid with similar viscosity (40% v/v glycerol/water solution with a viscosity of ~ 4 mPa·s) was also measured as a function of increasing pressure. As soon as pressure was applied (initial 1 kPa), the Newtonian reference started flowing, with flow increasing linearly with applied pressure. Notably, when pressure was removed, the Newtonian reference continued flowing out of the nozzle. The two venom mimics with a similar *η*_∞_ (*mimic I* with 200 mg/mL BSA protein and *mimic II* with 1 mg/mL xanthan gum) were also measured as a function of increasing pressure. Similar to the real venom, they only exhibited flow after a pressure threshold was reached. In the 0-3 kPa regime no flow was observed, in the 3-7 kPa regime discontinuous dripping was observed, and in the >7 kPa regime continuous flow was observed (**Figure 6b**). After the initial no-flow regime, both mimic formulations exhibited a linear flow trend with increasing pressure, overlapping with the behavior observed in real venom samples and indicating a good match of the shear-thinning properties. To further corroborate the shear-thinning behavior and controllable delivery of fluid, cyclic measurements were performed turning the pressure on and off at varying pressures (**Figure 6c**). When the pressure was turned off, the venom mimic fluid immediately stopped flowing with no visible dripping, and the same flow was recovered when the pressure was reapplied. To demonstrate this control, we incorporated a dye into the *venom mimic II* formulation and deposited droplets with controlled volume at specified locations by turning on/off the pressure while the nozzle moved along a preprogrammed path (**Figure 6d**). This switching flow behavior provides good control over the venom delivery mechanisms, and therefore can be used to delivery fluid at specific flow rates at desired times by controlling pressure.

**Figure 6.**
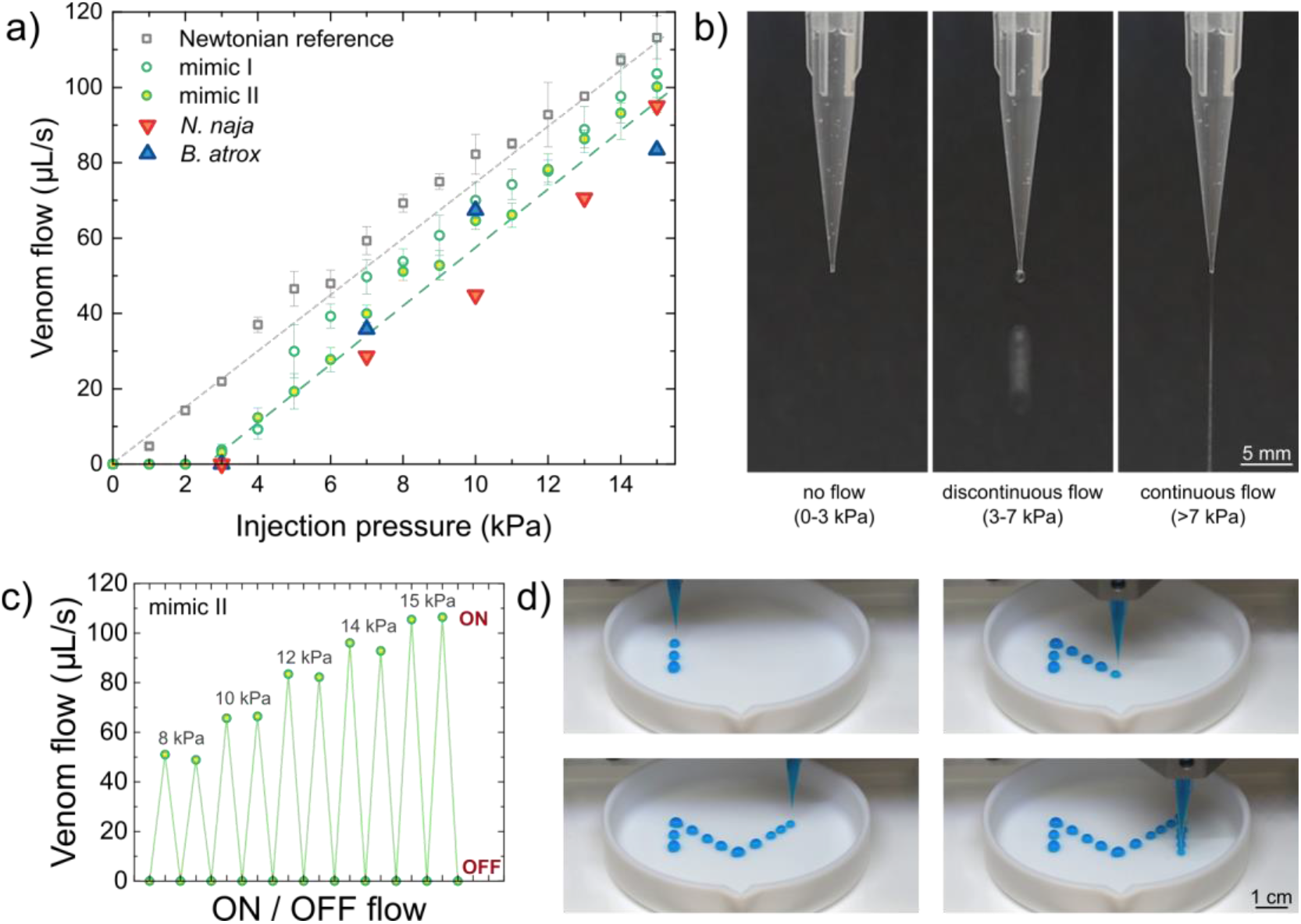
Simulated venom delivery in a direct ink writing printing system. **a)** Venom flow as a function of applied pressure. Real venom (*N. naja* and *B. atrox*) and *venom mimics I* and *II* exhibited flow only after a minimum pressure threshold was achieved. **b)** Venom mimics exhibited a no-flow regime (0-3 kPa), a discontinuous dripping flow regime (3-7 kPa), and a continuous flow regime (>7 kPa). **c)** Venom mimics exhibited switchable flow when the pressure was turned on or off. **d)** The controlled delivery of venom mimics was demonstrated by direct ink writing of a colored mimic II sample over a preprogrammed “M” path.

## Conclusions

We have analyzed the rheological properties of venom from thirteen snake species (seven vipers and six elapids) and studied how their viscosity is affected by shear rate. The venom exhibited shear-thinning behavior over the range of study, showing lower viscosities at high rates. We modeled this non-Newtonian behavior with an Ostwald–de Waele power-law, where all analyzed venom samples consistently exhibited shear-thinning properties. This flow behavior would aid the envenomation process, where high viscosity at rest would help to store the venom in the glands and low viscosity under pressure (causing high shear rates) would help flow during delivery. However, we did not find evidence for phylogenetic signal in the measured venom viscosities, suggesting that rheological parameters of venom may vary independently of relationships between snake species.

In addition, this viscosity characterization study provided useful data to design venom mimic surrogate fluids that can reproduce the rheological properties of venom in inexpensive, safe formulations. Protein-based mimics consisting of BSA aqueous solutions spanning concentrations up to 200 mg/mL showed poor performance as venom mimics, as the measured viscosities at low shear rates was significantly lower than real venom samples. Venom mimics based on xanthan gum, on the other hand, exhibited much higher viscosities over the studied rates, providing a broad range of flow properties and shear-thinning behaviors as a function of concentrations (up to 3 mg/mL). This second mimic system spans flow properties within the range of real venom and can be formulated to reproduce specific flow behaviors characteristic of individual snake species.

We further evaluated the performance of these venom mimic surrogate fluids in a simulated venom delivery system using a DIW 3D printer (which can simulate the extrusion of shear-thinning biofluids through a microchannel with controlled pressure). In both real and mimic venoms, we observed a three-regime flow behavior as a function of pressure. At low pressures (0-3 kPa) no flow was observed (rest state), at intermediate pressures (0-3 kPa) venom was dripping discontinuously, and above a threshold pressure (>7 kPa) continuous flow was observed. This on/off flow behavior was observed over multiple delivery cycles, providing good control over the venom delivery mechanism by controlling pressure.

Although this simulated venom delivery system has been designed to conceptually model the extrusion of venom through the snake fangs, real envenomation is of course much more complex. Snake fangs have smaller microchannels and more complex structures, combining different morphological features that contribute to other predatory functions as well (e.g., puncture). However, it provides a robust testbed to evaluate and quantitatively compare the different venoms, to guide the design of surrogate venom mimics, and to study envenomation physics in an inexpensive and safe method. Building upon the work described here, it is possible to design species-specific venom mimics that facilitate further research on the biomechanics and fluid dynamics of venom delivery *via* snake bites, improve snakebite emergency response treatment protocols, optimize protective clothing design, and inform the design of bioinspired puncture and injection devices.

## Acknowledgements

All authors acknowledge the support of the University of Michigan College of Engineering for partial support of this research.

## Author contributions

M.L.H., T.Y.M., and A.P.F. conceived, designed, and supervised the research. M.F. prepared all venom samples and performed the rheological characterization experiments. M.L.H. performed electrophoretic and phylogenetic comparative analyses. Y.L. performed the simulated delivery tests. All authors participated in manuscript revisions, discussions, and interpretation of the data.

## Competing interests

The authors declare no competing interests.

## Data and materials availability

All data is available in the main text or in the supporting materials.

## Notes

### Competing Interest Statement

The authors have declared no competing interest.

## References

(1) Seifert, S. A.; Armitage, J. O.; Sanchez, E. E. Snake Envenomation. N Engl J Med 2022, 386 (1), 68–78. 10.1056/NEJMra2105228.

(2) Vázquez Torres, S.; Benard Valle, M.; Mackessy, S. P.; Menzies, S. K.; Casewell, N. R.; Ahmadi, S.; Burlet, N. J.; Muratspahić, E.; Sappington, I.; Overath, M. D.; Rivera-de-Torre, E.; Ledergerber, J.; Laustsen, A. H.; Boddum, K.; Bera, A. K.; Kang, A.; Brackenbrough, E.; Cardoso, I. A.; Crittenden, E. P.; Edge, R. J.; Decarreau, J.; Ragotte, R. J.; Pillai, A. S.; Abedi, M.; Han, H. L.; Gerben, S. R.; Murray, A.; Skotheim, R.; Stuart, L.; Stewart, L.; Fryer, T. J. A.; Jenkins, T. P.; Baker, D. De Novo Designed Proteins Neutralize Lethal Snake Venom Toxins. Nature 2025. 10.1038/s41586-024-08393-x.

(3) Oliveira, A. L.; Viegas, M. F.; Da Silva, S. L.; Soares, A. M.; Ramos, M. J.; Fernandes, P. A. The Chemistry of Snake Venom and Its Medicinal Potential. Nat Rev Chem 2022, 6 (7), 451–469. 10.1038/s41570-022-00393-7.

(4) Sánchez, E. E.; Galán, J. A.; Powell, R. L.; Reyes, S. R.; Soto, J. G.; Russell, W. K.; Russell, D. H.; Pérez, J. C. Disintegrin, Hemorrhagic, and Proteolytic Activities of Mohave Rattlesnake, Crotalus Scutulatus Scutulatus Venoms Lacking Mojave Toxin. Comparative Biochemistry and Physiology Part C: Toxicology & Pharmacology 2005, 141 (2), 124–132. 10.1016/j.cca.2005.04.001.

(5) Tasoulis, T.; Isbister, G. A Review and Database of Snake Venom Proteomes. Toxins 2017, 9 (9), 290. 10.3390/toxins9090290.

(6) Kardong, K. V.; Lavin-Murcio, P. A. Venom Delivery of Snakes as High-Pressure and Low-Pressure Systems. Copeia 1993, 1993 (3), 644–650. 10.2307/1447225.

(7) Cundall, D. Viper Fangs: Functional Limitations of Extreme Teeth. Physiological and Biochemical Zoology 2009, 82 (1), 63–79. 10.1086/594380.

(8) Ewoldt, R. H.; Saengow, C. Designing Complex Fluids. Annu. Rev. Fluid Mech. 2022, 54 (1), 413–441. 10.1146/annurev-fluid-031821-104935.

(9) Triep, M.; Hess, D.; Chaves, H.; Brücker, C.; Balmert, A.; Westhoff, G.; Bleckmann, H. 3D Flow in the Venom Channel of a Spitting Cobra: Do the Ridges in the Fangs Act as Fluid Guide Vanes? PLOS ONE 2013, 8 (5), e61548. 10.1371/journal.pone.0061548.

(10) Challita, E. J.; Rohilla, P.; Bhamla, M. S. Fluid Ejections in Nature. Annual Review of Chemical and Biomolecular Engineering 2024, 15 (Volume 15, 2024), 187–217. 10.1146/annurev-chembioeng-100722-113148.

(11) Cleuren, S. G. C.; Hocking, D. P.; Evans, A. R. Fang Evolution in Venomous Snakes: Adaptation of 3D Tooth Shape to the Biomechanical Properties of Their Prey. Evolution 2021, 75 (6), 1377–1394. 10.1111/evo.14239.

(12) Young, B. A.; Herzog, F.; Friedel, P.; Rammensee, S.; Bausch, A.; Van Hemmen, J. L. Tears of Venom: Hydrodynamics of Reptilian Envenomation. Phys. Rev. Lett. 2011, 106 (19), 198103. 10.1103/PhysRevLett.106.198103.

(13) Mackessy, S. P. Venom Composition in Rattlesnakes: Trends and Biological Significance. In The biology of rattlesnakes; 2008.

(14) de Alencar, L. R. V.; Martins, M.; Burin, G.; Quental, T. B. Arboreality Constrains Morphological Evolution but Not Species Diversification in Vipers. Proceedings of the Royal Society B: Biological Sciences 2017, 284 (1869), 20171775. 10.1098/rspb.2017.1775.

(15) Holding, M. L.; Trevine, V. C.; Zinenko, O.; Strickland, J. L.; Rautsaw, R. M.; Mason, A. J.; Hogan, M. P.; Parkinson, C. L.; Grazziotin, F. G.; Santana, S. E.; Davis, M. A.; Rokyta, D. R. Evolutionary Allometry and Ecological Correlates of Fang Length Evolution in Vipers. Proceedings of the Royal Society B: Biological Sciences 2022, 289 (1982), 20221132. 10.1098/rspb.2022.1132.

(16) Avella, I.; Barajas-Ledesma, E.; Casewell, N. R.; Harrison, R. A.; Rowley, P. D.; Crittenden, E.; Wüster, W.; Castiglia, R.; Holland, C.; Van Der Meijden, A. Unexpected Lack of Specialisation in the Flow Properties of Spitting Cobra Venom. Journal of Experimental Biology 2021, 224 (7), jeb229229. 10.1242/jeb.229229.

(17) Smith, P. K.; Krohn, R. I.; Hermanson, G. T.; Mallia, A. K.; Gartner, F. H.; Provenzano, M. D.; Fujimoto, E. K.; Goeke, N. M.; Olson, B. J.; Klenk, D. C. Measurement of Protein Using Bicinchoninic Acid. Analytical Biochemistry 1985, 150 (1), 76–85. 10.1016/0003-2697(85)90442-7.

(18) Revell, L. J. Phytools: An R Package for Phylogenetic Comparative Biology (and Other Things). Methods Ecol Evol 2012, 3 (2), 217–223. 10.1111/j.2041-210X.2011.00169.x.

(19) Kumar, S.; Suleski, M.; Craig, J. M.; Kasprowicz, A. E.; Sanderford, M.; Li, M.; Stecher, G.; Hedges, S. B. TimeTree 5: An Expanded Resource for Species Divergence Times. Molecular Biology and Evolution 2022, 39 (8), msac174. 10.1093/molbev/msac174.

(20) Venomous Reptiles and Their Toxins: Evolution, Pathophysiology, and Biodiscovery; Fry, B., Ed.; Oxford University Press: New York, NY, 2015.

(21) Utkin, Y. N. Last Decade Update for Three-Finger Toxins: Newly Emerging Structures and Biological Activities. WJBC 2019, 10 (1), 17–27. 10.4331/wjbc.v10.i1.17.

(22) Olaoba, O. T.; Karina Dos Santos, P.; Selistre-de-Araujo, H. S.; Ferreira De Souza, D. H. Snake Venom Metalloproteinases (SVMPs): A Structure-Function Update. Toxicon: X 2020, 7, 100052. 10.1016/j.toxcx.2020.100052.

(23) Markland, F. S.; Swenson, S. Snake Venom Metalloproteinases. Toxicon 2013, 62, 3–18. 10.1016/j.toxicon.2012.09.004.

(24) Cerda, P. A.; Crowe-Riddell, J. M.; Gonçalves, D. J. P.; Larson, D. A.; Duda, T. F.; Davis Rabosky, A. R. Divergent Specialization of Simple Venom Gene Profiles among Rear-Fanged Snake Genera (Helicops and Leptodeira, Dipsadinae, Colubridae). Toxins 2022, 14 (7), 489. 10.3390/toxins14070489.

(25) Larson, R. G. The Structure and Rheology of Complex Fluids; Topics in chemical engineering; Oxford university press: New York, 1999.

(26) Aghakhani, A.; Pena-Francesch, A.; Bozuyuk, U.; Cetin, H.; Wrede, P.; Sitti, M. High Shear Rate Propulsion of Acoustic Microrobots in Complex Biological Fluids. Sci. Adv. 2022, 8 (10), eabm5126. 10.1126/sciadv.abm5126.

(27) Young, B. A.; Herzog, F.; Friedel, P.; Rammensee, S.; Bausch, A.; Van Hemmen, J. L. Tears of Venom: Hydrodynamics of Reptilian Envenomation. Phys. Rev. Lett. 2011, 106 (19), 198103. 10.1103/PhysRevLett.106.198103.

(28) Triep, M.; Hess, D.; Chaves, H.; Brücker, C.; Balmert, A.; Westhoff, G.; Bleckmann, H. 3D Flow in the Venom Channel of a Spitting Cobra: Do the Ridges in the Fangs Act as Fluid Guide Vanes? PLoS ONE 2013, 8 (5), e61548. 10.1371/journal.pone.0061548.

(29) Ikeda, S.; Nishinari, K. Intermolecular Forces in Bovine Serum Albumin Solutions Exhibiting Solidlike Mechanical Behaviors. Biomacromolecules 2000, 1 (4), 757–763. 10.1021/bm005587o.

(30) Zhang, Z.; Liu, Y. Recent Progresses of Understanding the Viscosity of Concentrated Protein Solutions. Current Opinion in Chemical Engineering 2017, 16, 48–55. 10.1016/j.coche.2017.04.001.

(31) Castellanos, M. M.; Pathak, J. A.; Colby, R. H. Both Protein Adsorption and Aggregation Contribute to Shear Yielding and Viscosity Increase in Protein Solutions. Soft Matter 2014, 10 (1), 122–131. 10.1039/C3SM51994E.

(32) Milas, M.; Rinaudo, M.; Knipper, M.; Schuppiser, J. L. Flow and Viscoelastic Properties of Xanthan Gum Solutions. Macromolecules 1990, 23 (9), 2506–2511. 10.1021/ma00211a018.

(33) Speers, R. A.; Tung, M. A. Concentration and Temperature Dependence of Flow Behavior of Xanthan Gum Dispersions. Journal of Food Science 1986, 51 (1), 96–98. 10.1111/j.1365-2621.1986.tb10844.x.

(34) Garcıá-Ochoa, F.; Santos, V. E.; Casas, J. A.; Gómez, E. Xanthan Gum: Production, Recovery, and Properties. Biotechnology Advances 2000, 18 (7), 549–579. 10.1016/S0734-9750(00)00050-1.

(35) Nsengiyumva, E. M.; Alexandridis, P. Xanthan Gum in Aqueous Solutions: Fundamentals and Applications. International Journal of Biological Macromolecules 2022, 216, 583–604. 10.1016/j.ijbiomac.2022.06.189.

